# Interpretable Machine Learning Uncovers Structural Determinants of Wnt-Wls Binding Specificity from Extended Atomistic Simulations

**DOI:** 10.1101/2025.08.18.670971

**Authors:** Tiffany J. Callahan, Jie Shi, Kevin J. Cheng, Michael A. Sauer, Taras V. Pogorelov, Sara Capponi

## Abstract

The Wnt protein family plays a critical role in cell development, with each Wnt protein interacting differently with the Wls membrane protein through distinct binding residues. A direct comparison and elucidation of the molecular mechanisms underlying Wnt–Wls binding across the diverse Wnt family remain challenging, owing to variations in sequence length and amino acid composition among Wnt proteins, which can affect their binding affinity and trafficking efficiency via Wls. Here we combine extended atomistic molecular dynamics simulations with supervised machine learning to elucidate binding specificity among four Wnt proteins, selected based on experimental structure availability and scientific relevance. We implement a local structure alignment algorithm to enable cross-system matching and comparison of residue interactions, and we apply a two-stage clustering strategy to reduce feature redundancy and facilitate robust feature selection. After training a Random Forest classifier that achieved high predicting accuracy, our feature importance analysis reveals both previously known and novel key residue pairs responsible for distinguishing among the Wnt systems. Our findings highlight that the binding specificity across different systems arises from the distributed nature of interactions across the protein binding surface and demonstrate how interpretable machine learning can effectively uncover crucial biophysical interactions. Importantly, our integrated strategy is generalizable to other systems and provides a data-driven approach for analyzing protein– protein interactions and guiding experimental validation or therapeutic targeting.

## 1. Introduction

The Wnt family of secreted glycoproteins plays a central role in embryonic development, tissue homeostasis, and the regulation of cell signaling across multicellular organisms^1–6^. Aberrant Wnt signaling is implicated in numerous human diseases^7^, including cancer^8–14^, neurodevelopmental and neurodegenerative disorders^15–17^, and bone diseases^18,19^. These diverse biological outcomes arise from the interaction of 19 human Wnt proteins with a wide array of receptors and co-receptors, enabling highly context-dependent signaling cascades^20–24^. Despite their functional diversity, all Wnt proteins rely on a single cargo receptor, Wntless (Wls), for secretion and delivery to the plasma membrane^21,25,26^. Wls is a conserved multi-pass membrane protein that binds Wnts within the secretory pathway, facilitating their trafficking and eventual release for extracellular signaling^26–28^. The fact that a structurally conserved receptor can bind to such a wide range of ligands raises a fundamental question: what molecular features allow it to recognize such diverse partners with specificity? Understanding this mechanism would clarify how Wnt signaling maintains fidelity and shed light on how mutations in Wnt or Wls can disrupt binding and contribute to disease. Answering this question requires methods capable of resolving the fine-grained, dynamic nature of protein-protein interactions while accommodating substantial sequence and conformational variability across Wnt family members^29–32^. Static structural data are insufficient, as Wnt-Wls binding involves flexible domains, lipid modifications, and dynamic contact networks^21,25,27^. To address these challenges, we combined atomistic molecular dynamics (MD) simulations with supervised machine learning (ML) to systematically identify the structural determinants of Wnt-Wls binding specificity.

Recent studies have demonstrated the promise of combining MD simulations and ML to dissect protein-protein interactions ^33–39^. Specifically relevant for the scope of our work were two studies focused on deciphering the contributions of each amino acid and of their mutations to the intermolecular interactions. Capponi et al. studied the interaction between the ACE2 human receptor and the SARS-CoV-2 binding domain using Convolutional Neural Networks (CNNs) trained on images generated from the protein-protein contact matrix computed from extended MD simulations of different SARS-CoV-2 mutants bound to ACE-2^29^. The authors accurately predicted binding affinity trends across SARS-CoV-2 spike protein mutants. In the same year, Pavlova et al. used a Random Forest (RF) classifier trained on MD-derived contact matrices to differentiate SARS-CoV and SARS-CoV-2 binding interactions, identifying key affinity-modulating residues^30^. The approach used in these two works illustrates how dynamic contact features and interpretable ML models can uncover mechanistic insights from MD simulation data. Inspired by these works, we sought to apply a similar strategy to a different biological problem: the recognition of a structurally conserved membrane protein (Wls) by a diverse protein family (Wnts). Wnt proteins differ substantially in both sequence length and composition, necessitating the development of new methods for aligning and comparing residue interactions across systems. To this end, we developed a novel, local structure alignment algorithm and a carefully design ML workflow to identify conserved contact features and infer determinants of Wnt-Wls specificity.

We focused on four Wnt proteins, Wnt1, Wnt3a, Wnt5a, and Wnt8a, selected based on structural availability and their scientific prominence. Wnt3a and Wnt8a were available in experimentally resolved forms^25,27^, while Wnt1 and Wnt5a were modeled via template-based homology modeling. Together, these systems span canonical and noncanonical signaling pathways and represent both conserved and divergent members of the Wnt family^40^. Each Wnt-Wls complex was simulated for 1.5 microseconds in two sets of independent simulations, yielding rich trajectory datasets for dynamic contact analysis. A central challenge in comparing these systems arises from differences in Wnt sequence length and composition. While it is standard practice to apply sequence alignment based on amino acid types, such an approach may not accurately reflect the true interactional relationships, as contact formation is inherently three-dimensional and closely tied to the protein’s structural context^30^. Furthermore, traditional sequence alignment methods may fail to identify structurally analogous residues when sequence identity is low. To address this, we developed a novel RMSD-based local structural algorithm to map residue contact pairs across all four Wnt-Wls complexes to a shared structural reference. This enabled us to define a consistent set of residue-residue contacts across all systems, preserving conserved interaction patterns while tolerating sequence divergence.

From the MD simulations, we extracted Wnt-Wls residue pairs within a 12 Å contact distance, resulting in a shared set of structurally aligned contact pairs across all complexes. Time series of these contact distances were used as features to train a RF classifier capable of distinguishing among the four Wnt proteins. To ensure statistical independence in our model evaluation, we developed a method based on autocorrelation function (ACF) for partitioning temporally uncorrelated data in training and test sets. Further dimensionality reduction via hierarchical clustering enabled selection of the most informative contact features for model training. The resulting RF classifier achieved a test accuracy of 95.9%, demonstrating the model’s ability to differentiate Wnt-Wls binding modes with high precision. Permutation feature importance analysis revealed key residue pairs located near known functional regions, including the palmitoleoylation (PAM) site and conserved hairpin motifs. Notably, several top-ranked features overlapped with residues previously implicated in Wnt secretion or Wls recognition through biochemical studies^16,17^, lending biological validation to our findings.

Beyond its relevance to Wnt biology, our approach offers a generalizable framework for analyzing protein-protein interactions among homologous ligands or receptors. The RMSD-based local structural alignment strategy facilitates structural comparisons across divergent family members, while our interpretable ML pipeline extracts functional signals from complex, time-resolved simulation data. Notably, our results show that even relatively simple ML models, such as RFs, can yield biologically meaningful insights when paired with high-quality simulations and domain-specific feature engineering. More broadly, this study demonstrates how integrating MD simulations and ML can reveal the molecular determinants of receptor recognition in systems characterized by structural diversity and functional redundancy. In the case of Wnt-Wls, this approach illuminates how specificity is achieved despite sequence variation, an insight with direct relevance for understanding how mutations disrupt Wnt signaling and contribute to disease. By bridging atomistic dynamics with predictive modeling, our framework provides a scalable and biologically grounded path toward understanding the determinants of protein-protein interactions in other systems of biomedical importance.

## 2. Results

### 2.1 Identification of Wnt-Wls Systems Based on Experimental Structures and Biological Significance

By interacting with different receptors, Wnt proteins regulate a variety of processes including cell proliferation, migration, survival, and polarity. Based on the pathways they activate, Wnt proteins has been traditionally categorized as canonical (β-catenin-dependent) or noncanonical (β-catenin-independent)^7^. However, this classification is not strict as both canonical and noncanonical Wnt signaling can be activated by individual Wnt proteins depending on specific cellular context^6,7^. For the scope of this work, we first identified Wnt3a and Wnt8a because the availability of the experimental structures^25,27^ allowed us to set up and perform extended atomistic simulations and build models for the other systems. Wnt3a and Wnt8a are both considered canonical pathway activators^7,41^. Then we selected Wnt1 and Wnt5a for their scientific importance. Wnt1 signaling pathway is categorized as canonical, while that of Wnt5a as noncanonical and is related with cell polarity and migration. All the selected systems are associated with different human diseases, and either their signaling pathways or alterations therein have been linked to different types of cancer^14,21,40^. To confirm the scientific relevance of our selection, we also considered different metrics such as total number of publications, citations, and H-index and reported them in Figure 1. Overall, Wnt3a emerges as the most extensively studied with 2,517 publications, 101,826 citations, and H-index of 144, and it is followed by Wnt5a with substantial research activity reflected by 2,387 publications and an H-index of 128. Wnt1 is similarly influential, ranking third with 1,872 publications and an H-index of 128.

**Figure 1.**
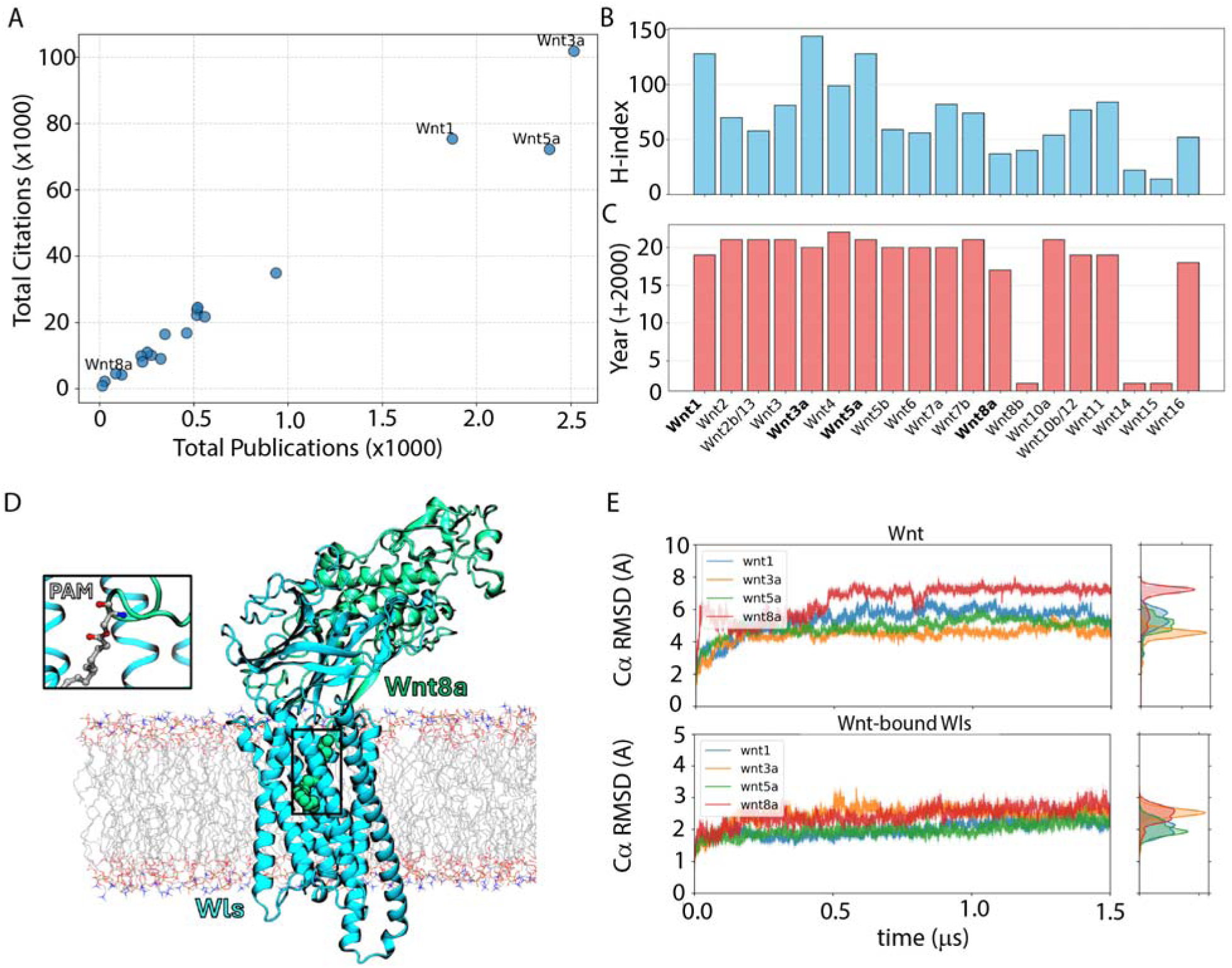
Comparative Bibliometric and Structural Characterization of Wnt-Wls Complexes. (A) Scatter plot showing the correlation between total publications and total citations per Wnt protein. (B) H-index values are reported as functions of Wnt proteins. (C) Peak year of research activity (by publication count) for each Wnt protein across the past decades from year 2000, based on data from the Web of Science Core Collection. (D) Structural snapshot from MD simulations showing Wnt8a (green) bound to Wls (cyan) within a membrane. The lipid bilayer is transparent, revealing Wls’ membrane-embedded conformation. The inset highlights the palmitoleic acid (PAM) lipid modification on Wnt8a, shown in licorice representation, inserting into the hydrophobic groove of the Wnt-Wls interface. (E) Time evolution of C_α_ RMSD during MD simulations for each Wnt protein (top) and their corresponding Wnt-bound Wls partners (bottom). Right-side kernel density estimates (KDEs) show RMSD distributions.

### 2.2 Revealing the Dynamics of Wnt-Wls Complexes through Extended MD Simulations

To perform the MD simulations, we used the initial coordinates of Wnt8a-Wls and Wnt3a-Wls complexes from the experimental structures PBD ID 7DRT^25^ and 7KC4 ^27^, respectively. For the Wnt1- and Wnt5a-Wls systems, we modeled the initial coordinates using Phyre2.2^42^ (see Methods). Then we constructed the forcefield parameters to describe the mono-unsaturated palmitoleate (PAM) covalently attached to a conserved serine residue present in the human Wnts (inset of Figure 1C and Methods). We used an approach based on building the forcefield parameter files by leveraging existing ones for similar molecules and we were able to describe Wnt-Wls interactions and capture PAM’s essential function, such as its anchoring to the lipid bilayer^27^. The four Wnt-Wls systems were embedded in a palmitoyloleoylphosphatidylcholine (POPC) bilayer in excess water and equilibrated. The production runs were carried out in *NPT* ensemble (constant number of particles *N*, pressure *P*, and temperature *T*) and the simulations were extended for 1.5 μs length for each system. We performed two independent sets of simulations for each system. In Figure 1C, we show the initial structure of the Wnt8a-Wls simulation as a representative of all the systems studied. Also, we used the Wnt8a-Wls complex as reference for the ML analysis.

To analyze the average dynamics of the systems, we computed the time evolution of the root-mean squared deviations (RMSDs) for the Wnt and Wls CL atoms and represented them in Figure 1D. The RMSD calculation provides a measure of the average motion of a protein from its initial structure at each simulated time step. Overall, the four Wnt proteins exhibit high mobility with RMSDs ranging from an average of approximately 4.5 Å for Wnt3a to an average of approximately 7Å for Wnt8a (Figure 1D, top panel). After an initial increase, we observe that the RMSD values reach a plateau after 500 ns. The Wls proteins appear to be more rigid with the CL atom RMSDs ranging from an average of approximately 2 Å for Wnt5a and Wnt1 to an average of approximately 2.5 Å for Wnt8a and Wnt3a. These RMSD low values indicate that the dynamics of the Wls proteins in the four systems is dominated by the transmembrane region that has limited motion being embedded in the bilayer. The RMSDs calculated from the Replica 2 simulations (Supplementary Material Figure S1) show values consistent with those observed in Replica 1. The RMSD values of the Wnt proteins span between 4 and 6 Å and those of Wls proteins between 1.6 and 2.9 Å. We then compared the RMSDs of each Wnt and Wls proteins of Replica 1 simulation with the corresponding protein of Replica 2 simulation (Supplementary Material Figure S2). Overall, the RMSD distributions of the two replicas reveal differences that are on average less than 2 Å, indicating that the protein dynamics are similar during the two independent simulation sets.

We then examined the local dynamical features by computing the root mean squared fluctuations (RMSFs), which measure the time-averaged fluctuations of an amino acid position during the simulations (see <u>Methods</u>). In Supplementary Material Figure S3, we represented the RMSF calculated over the last 500 ns of the simulations for the Wnt and Wls proteins for Replica 1 and 2. From a quick look at the results, we observe high similarity in the RMSF patterns between the two replicas with the majority of the residues showing significantly different RMSF values located far away from the protein binding interface. Because our work is focused on the Wnt-Wls contact area, we assume that these residues do not affect our analysis. To better investigate the RMSF difference between the two replicas and ensure the identification of the residues, if any, exhibiting distinct local dynamics, we used a dual filter and analyzed every single residue between Replica 1 and 2 for all systems. We then employed a two-tailed Student’s t-test with Benjamini-Hochberg (BH) correction to control the false discovery rate to perform the statistical comparison and studied only the residues with mean RMSF difference greater than 1 Å between the two replicas. We found that all residues of Wnt3a, Wnt-5a, and Wnt-8a passing the filter are far away from the binding region, whereas residues R156 and T231 in Wnt1 located in proximity of the binding interface exhibit a mean RMSF difference of 1.01 and 1.89 Å respectively. Based on our strategy, these two residues are the only ones that would warrant further investigation in case they are identified as determinants for the Wnt-Wls interactions. Given that the RMSD and RMSF analysis show that the two independent sets of simulations show highly similar dynamics, we performed the downstream analysis only on Replica 1.

#### Local Structure Alignment

After analyzing the dynamics of the Wnt and Wls proteins in the four systems under investigation, we shift our focus to the contact interface. This analysis presents a unique challenge: how to define a comparison strategy across different Wnt proteins within the same family? Unlike previous studies that examine a single protein or protein complex under varying binding modes or conformational states^29,39^, our work involves four distinct Wnt proteins that differ in both sequence length and amino acid composition. While traditional approaches that define contact regions based on distance cutoffs^29,30^ are appropriate for analyzing individual systems, they are insufficient for identifying conserved contact pairs across Wnt proteins due to their sequence divergence. A standard approach, as employed by Pavlova et al. in their comparison of SARS-CoV and SARS-CoV-2, involves performing sequence alignment based on amino acid types^30^. However, this method does not fully leverage structural information during alignment, potentially overlooking critical three-dimensional interaction features. To address this, we developed a structural alignment strategy to map residues across the four Wnt proteins and identify structurally analogous contact pairs. This approach allows us to construct a unified feature space, enabling direct comparison and downstream ML analysis. By aligning contact pair distances in this manner, we ensure that variations within each feature reflect meaningful differences in binding mode across Wnt isoforms.

To identify conserved Wnt-Wls contact pairs across the Wnt protein family, we developed an RMSD-based structural alignment algorithm (Figure 2A). Wnt8a was selected as the reference structure for this alignment, because it has the longer sequence of the two available experimentally resolved systems (at the time of analysis). This increases the likelihood that residues from the other Wnt proteins can be consistently mapped onto the reference, maximizing structural coverage and alignment fidelity across the datasets. Briefly, for each residue *i* in Wnt1, Wnt3a, or Wnt5a, we extracted a 41-residue segment centered in *i*, i. e. from *i*-20 to *i*+20. This segment was then structurally aligned to all possible 41-residue segments in Wnt8a, and the RMSD was calculated for each alignment. The residue *j* in Wnt8a that produced the lowest RMSD was identified as the best structural match to residue *i*. In Figure 2B we present the distribution of the lowest RMSD values for 41-residue segments from each residue of Wnt1, Wnt3a, and Wnt5a, obtained through structural alignment and mapping to Wnt8a. This process generated a residue-to-residue mapping dictionary for each Wnt protein relative to Wnt8a (the complete residue mapping dictionary is available in the GitHub repository). A more detailed description of the local alignment procedure and mapping strategy is provided in the <u>Methods section</u>.

**Figure 2.**
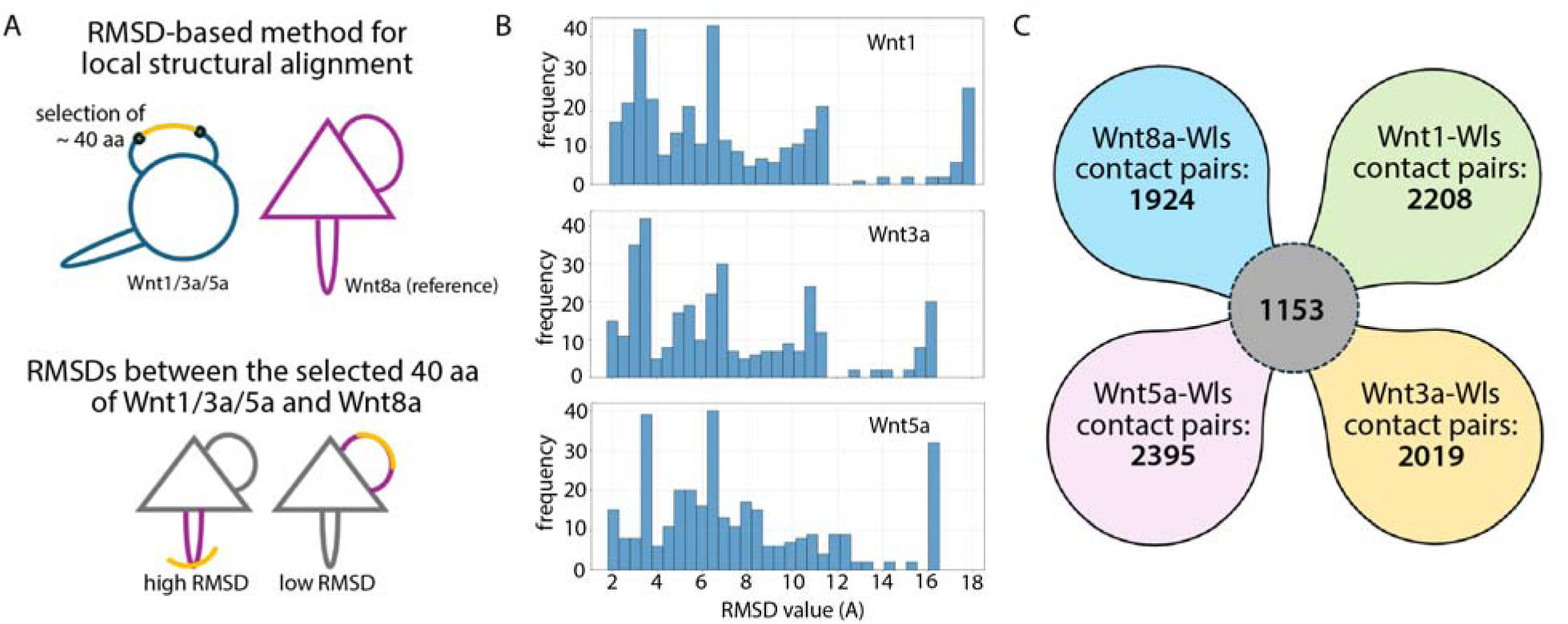
RMSD-Based Local Structural Alignment of Wnt Proteins to Identify Common Wnt-Wls Contact Pairs. (A) Scheme of the method for the structure-based alignments of the different Wnt sequences using the RMSDs and Wnt8a as reference. (B) Distribution of the lowest RMSD values calculated from local structure-based alignment for each residue between Wnt1 (top panel), Wnt3a (central panel), and Wnt5a (bottom panel) mapped onto Wnt8a. The x-axis represents the RMSD values and the y-axis indicates the frequency of each value observed. (C) Venn-like diagram showing the overlap of residue contact pairs identified across Wnt1, Wnt3a, Wnt5a, and Wnt8a. Each circle represents the total number of residue contact pairs unique to or shared among the Wnts. After mapping all the contact pairs to the reference structure Wnt8a, a total of 1,153 conserved contact pairs are present across all four Wnts as shown in the central intersection.

To ensure mapping quality, we applied a 10 Å RMSD threshold to determine alignment validity. Residues whose lowest RMSD value alignment exceeded this cutoff were considered to have no reliable structural counterpart in Wnt8a and were excluded from further analysis. This alignment-based approach enabled the identification of structurally conserved contact pairs across the four Wnt-Wls complexes, forming the basis for comparative analysis and feature extraction in downstream modeling.

Since the Wls protein maintains consistent residue indexing across all four systems, only the residue indices of Wnt1, Wnt3a, and Wnt5a were mapped to those of Wnt8a using the previously generated residue mapping dictionaries. This alignment yielded 1,153 structurally conserved contact pairs shared across all four Wnt-Wls complexes (Figure 2C, gray circle). Only these common contact pairs were retained as input features for subsequent data preprocessing for ML analysis.

Importantly, many of the aligned residue pairs involve different amino acid types. For example, E213 in Wnt3a aligns closely with S190 in Wnt8a, with an RMSD of approximately 1.93 Å, indicating strong structural similarity despite sequence divergence. This supports the notion that such substitutions can preserve local structure and should be retained for functional comparison, particularly in the context of Wls binding. Our alignment algorithm is specifically designed to identify these structurally conserved yet sequence-divergent residue pairs. These aligned residues were then used to identify Wnt-Wls contact pairs that are structurally comparable across all four systems.

Finally, we used a distance-based criterion to estimate the contact pairs for each individual Wnt-Wls complex. Two residues, one from Wnt and one from Wls, were considered to be in contact if their CL-CL distance was less than 12 Å in at least one frame of the simulation (see <u>Methods</u>). While alternative criteria have been applied in other studies^29,30^, we selected the 12 Å threshold based on the long-range interaction cutoff defined in the MD simulations. This choice ensures a comprehensive capture of all relevant Wnt-Wls contacts while maintaining a manageable dataset size for ML analysis. The resulting numbers of contact pairs were as follows: 2,208 for Wnt1-Wls, 2,019 for Wnt3a-Wls, 2,395 for Wnt5a-Wls, and 1,924 for Wnt8a-Wls (Figure 2C).

### 2.3 Unveiling key contact pairs in Wnt-Wls interactions that distinguish Wnt proteins using Machine Learning models

#### Autocorrelation-Informed Data Splitting for Model Training and Evaluation

Next we applied ML approaches to extract features from extended MD simulations (Figure 3). Atomistic simulations generate large datasets containing the Cartesian coordinates of every atom of each system at every time step. Due to the short simulation time step, the resulting trajectory frames are highly correlated, which poses a risk of data leakage and overfitting in ML models. To mitigate this, we developed an ACF-based method to partition the data into training and testing sets. Specifically, we computed the ACFs for each contact distance trajectory and averaged these across the 1,153 shared features for each Wnt-Wls system. We defined the minimum temporal gap for separating training and test data as the time τ at which the ACF decayed to 0 from its initial value. This ensured that frames used for testing were sufficiently decorrelated from those used for training. The estimated relaxation times were 332.6 ns for Wnt1, 379.04 ns for Wnt3a, 313.54 ns for Wnt5a, and 354.46 ns for Wnt8a, corresponding to 16,630, 18,952, 15,677, and 17,723 frames, respectively (see Supplementary Material Figure S4). Applying this method resulted in 162,036 training frames (Wnt1: 41,740; Wnt3a: 37,096; Wnt5a: 43,646; Wnt8a: 39,554) and 68,982 test frames (Wnt1: 16,630; Wnt3a: 18,952; Wnt5a: 15,677; Wnt8a: 17,723). The full procedure for ACF-based partitioning is detailed in the <u>Methods</u>.

**Figure 3.**
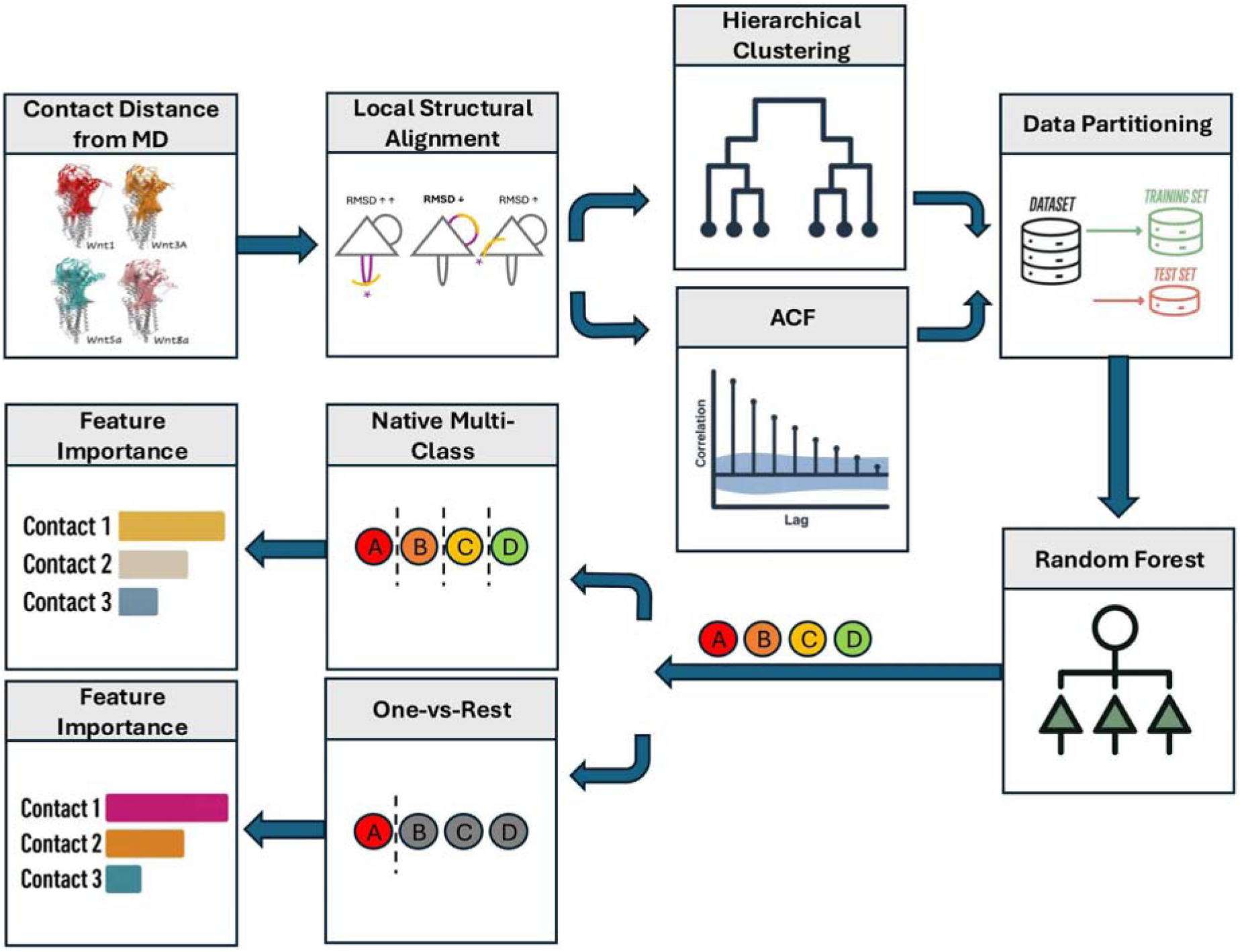
Workflow for Machine Learning-Driven Classification of Wnt-Wls Interaction Dynamics. Contact distances were extracted from MD simulations of four Wnt-Wls complexes. Local structural alignment was performed to identify common contact pairs for comparative analysis, followed by hierarchical clustering and ACF analysis to identify redundant features, and partition data into training and testing set to minimize temporal correlation. Relevant contact features from the training data set were used to train Random Forest classifiers. Two classification strategies were evaluated: native multi-class and one-vs-rest. Permutation feature importance was assessed to identify key contact pairs that distinguish between the four Wnt-Wls systems based on their dynamic contact profiles.

#### Identification of Common Wnt-Wls Contact Pairs via Hierarchical Clustering

After establishing a rigorous method to split the data in training and test sets that prevents data leakage, we performed feature selection using a two-round hierarchical clustering procedure aimed at reducing the number of input features and minimizing multicollinearity (Figure 4). In the first round, hierarchical clustering was applied to the 1,153 conserved contact pairs identified through local structural alignment. Residue indices were mapped to the Wnt8a reference system and grouped into five experimentally validated regions of interest^27^ (Figure 4C): hairpins 1 to 3, the N-terminus, and the palmitoleoylation (PAM) site.

**Figure 4.**
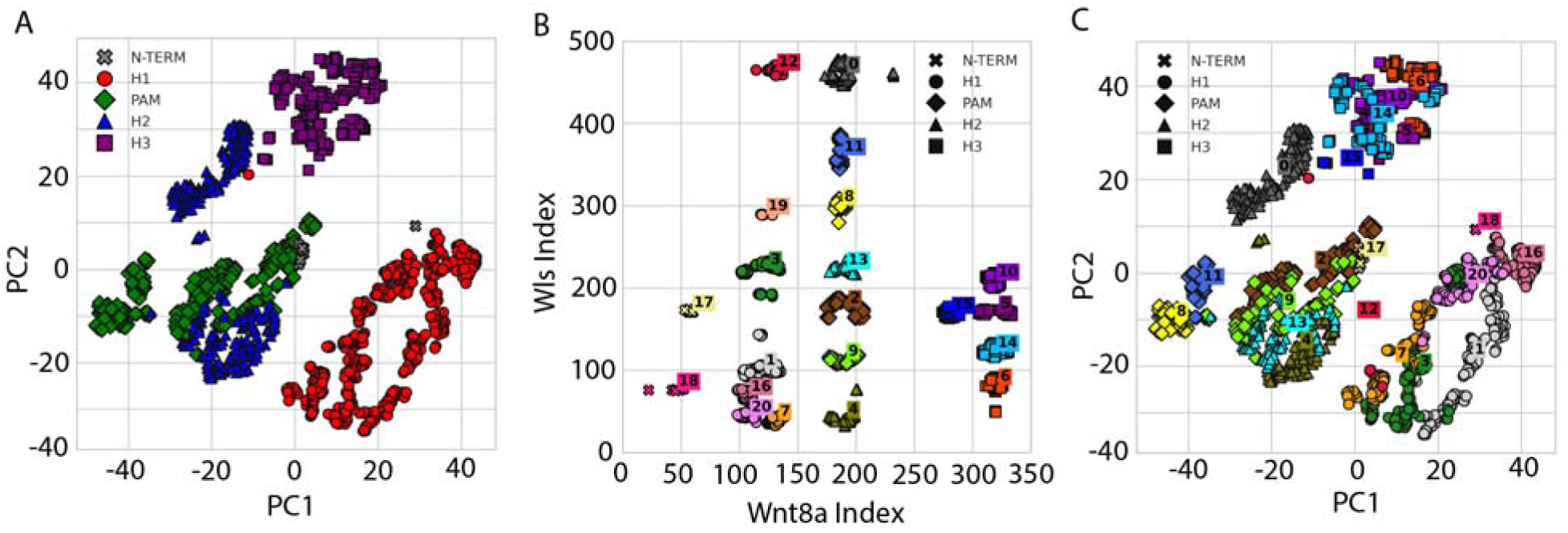
Clustering of Wnt8a Contact Pairs from Functionally Important Regions. A) t-SNE projection of Wnt8A contact pairs, colored by experimentally validated regions of interest: hairpins 1-3, the N-terminus, and the palmitoleoylation (PAM) site. B) hierarchical clustering of the Wnt8A contact pairs, revealing 21 clusters identified with the highest Silhouette score. Each cluster is represented with distinct colors and shapes corresponding to structural regions. C) illustrates the same t-SNE projection of contact distance pairs as (A), now colored by cluster assignment as (B). The 200 refined contact pairs represent features shared across the Wnts, optimized to reduce multicollinearity while retaining critical structural and functional information.

Using Silhouette analysis, we identified 21 primary clusters based on the maximum Silhouette score (SS = 0.76) (Figure 4B). For each cluster containing more than four features, a second round of hierarchical clustering was performed (<u>Methods</u>). This subclustering step yielded 200 subclusters, with an average SS of 0.40 (range: 0.24 to 1.0). From these subclusters, we selected one representative feature per cluster, resulting in a final set of 200 contact pairs used for model training and analysis (Figure 4A; Supplemental Table S1).

#### Determinants of Wnt-Wls Interactions Identified by Random Forest Classification

We trained an RF classifier using the 200 features previously identified. Following hyperparameter optimization using time series cross-validation on the training set, the optimal model configuration included 200 trees, a maximum tree depth of 3, and a minimum of 1,641 samples per leaf (<u>Methods</u>). Among several configurations that achieved perfect accuracy (100%) on both training and validation data, we selected the simplest model to minimize overfitting. To further assess model reliability, we generated a learning curve using a 20-fold time series split, with the relaxation time (τ) applied as the temporal gap between training and testing blocks. Accuracy consistently converged to 1.0 as the training size increased (Supplementary Material Figure S5), consistent with the results observed during hyperparameter tuning. Using the selected configuration, we retrained the model on the full training dataset. The model achieved 100% accuracy on the training set and 88.47% on the held-out test set. This performance drop reflects the effectiveness of our ACF-based partitioning strategy in minimizing temporal correlation, thereby allowing for a more realistic evaluation of generalization.

The modest decline in test accuracy was expected, as the testing set comprises frames from the early portion of the simulation trajectory (0 to τ), where structural fluctuations and contact variability are highest, as reflected in the RMSD profiles (Figure 1). These frames were excluded from all training and validation procedures and provide an independent, dynamic, and noise-prone region for evaluating model robustness. This setup ensures that the subsequent feature importance analysis is more likely to identify contact pairs that are not only discriminative but also resilient to structural variation across simulation conditions.

#### Feature Importance-Based Identification of Conserved Wnt-Wls Contacts

After establishing a model with high accuracy on both the train and test sets, we performed a feature importance analysis to identify the key contact features that differentiate interactions between the four Wnt-Wls systems. This analysis employed permutation importance, which quantifies each feature’s contribution by measuring the decrease in model accuracy when feature’s values are randomly shuffled. Because our model is a unified multiclass RF classifier, the resulting importance scores reflect each feature’s overall contribution to distinguishing among all four systems. To complement this global analysis with system-specific insights, we trained an additional one-vs-rest (OVR) classifier, in which a separate binary classifier is constructed for each Wnt-Wls system against the remaining three. Permutation importance was computed independently for each binary classifier, enabling the identification of features that are particularly informative for distinguishing a single system. This combined strategy allowed us to characterize both shared and system-specific determinants of Wnt-Wls binding interactions.

As shown in Figure 5A, the top contact pairs with the highest feature importance scores were identified using permutation analysis of the RF model. Overall, the absolute values of these scores are relatively small, with the highest reaching 0.0498. This is expected in models with a large number of features—such as the 200-feature model used in this study—where no single feature dominates the model’s predictive performance. Moreover, because permutation importance evaluates each feature independently by measuring its marginal effect while keeping all others fixed, it may underestimate the impact of features that are correlated or that contribute meaningfully only in combination with others. Nevertheless, in a high-performing model with low error rates, even a small absolute importance score can represent a substantial contribution to predictive accuracy.

**Figure 5.**
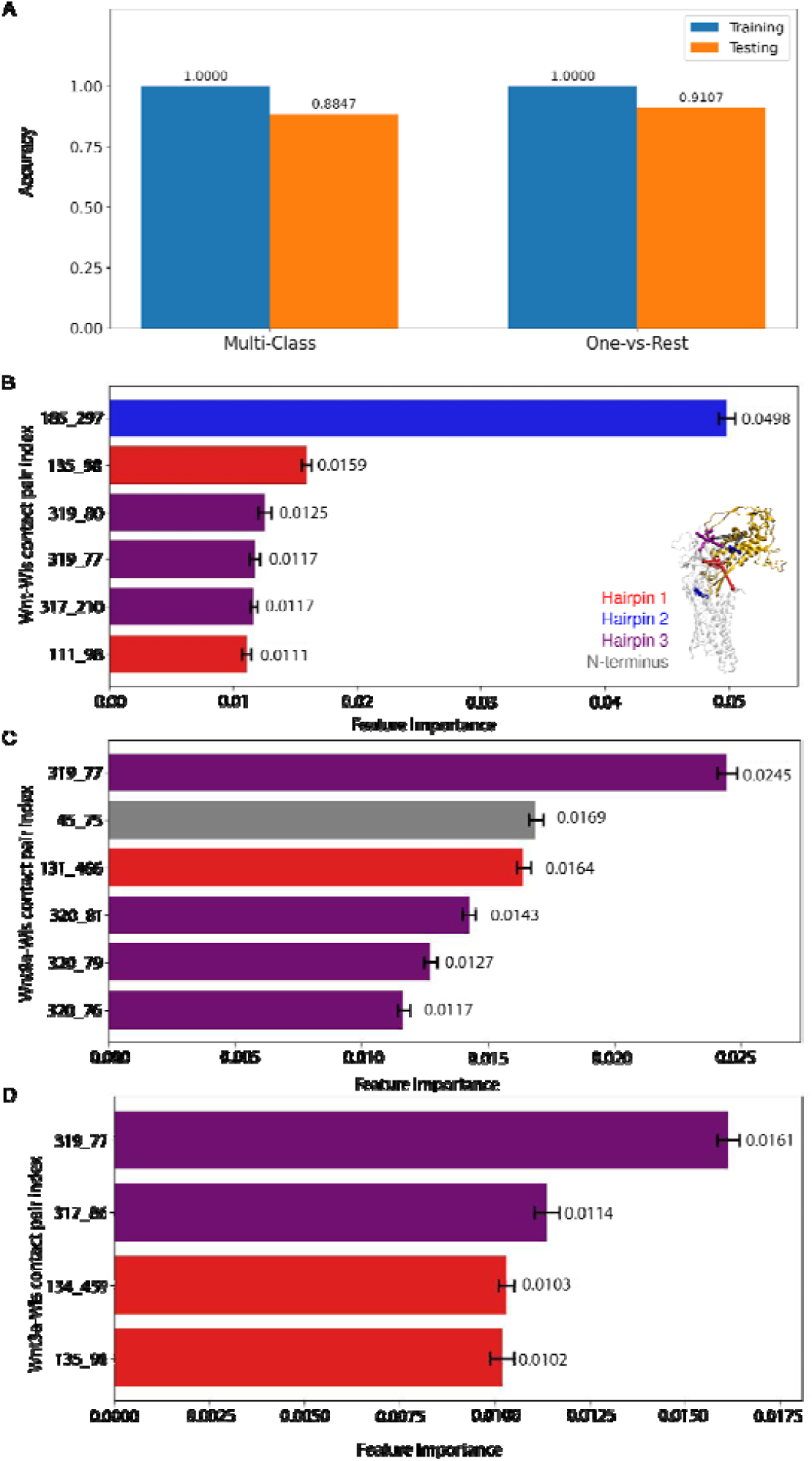
Classifier Performance and Key Features Distinguishing Wnt-Wls Complexes. (A) Accuracy comparison between native multi-class and one-vs-rest Random Forest classifiers, showing testing accuracy of 88.5% and 91.1%, respectively, with perfect training accuracy in both cases. (B-D) Permutation feature importance scores. The x-axis represents feature importance values, and the y-axis lists the top Wnt8aWnt-Wls contact pair indices. Error bars indicate variability across 100 iterations for each feature importance calculation. Color coding corresponds to structural elements in Wnt8a: red (Hairpin 1), blue (Hairpin 2), purple (Hairpin 3), and gray (N-terminus), as indicated by the structure illustration in inset of (B). (B) Top contact features contributing to the native multi-class classification. (C) and (D) Top contact features contributing to the one-vs-rest for Wnt 8a (C) and Wnt3a (D) respectively.

The contact pair I185-L297 was identified as the most influential feature in the classifier, with a feature importance score more than twice that of the next highest-ranked pair (Figure 5B). Notably, residue I185 is located adjacent to the PAM site in Wnt8a. Based on our local structural alignment and residue mapping dictionary, the corresponding residues in the other Wnt proteins are M223 in Wnt1, L208 in Wnt3a, and V243 in Wnt5a. Despite differences in amino acid identity, all of these residues occupy the same relative position next to the PAM site. This observation highlights the effectiveness of our alignment algorithm in capturing structurally analogous residues across divergent Wnt proteins. The PAM moiety is known to insert into the hydrophobic cavity of Wls and plays a critical role in Wnt-Wls binding. This site is highly conserved across nearly all Wnt isoforms, underscoring its essential role in Wnt biology^44^. The positioning of I185 and its counterparts in other Wnts suggests that these residues may facilitate the structural recognition or binding of Wnt proteins to Wls, thereby contributing to interaction specificity.

The contact pairs C319-A80 and C319-I77 ranked third and fourth in importance (Figure 5B). The corresponding residues for C319 across Wnt1, Wnt3a, and Wnt5a are C352, C334, and C362, respectively. These residues are identical in sequence and are located at the central position of the highly conserved W-C-C motif, which is evolutionarily preserved across the Wnt family^27^. Structural studies on Wnt3a have shown that C334 is critical for interaction with the hydrophobic pocket of Wls, and mutation of the equivalent residue in Wnt8a (C319) has been reported to abolish its signaling activity. These results support the functional significance of this motif and suggest that C319 and its structurally equivalent counterparts contribute to both Wls binding and downstream signaling.

#### System-Specific Feature Analysis Using One-vs-Rest Random Forests

To gain system-specific insights, we incorporated a OneVsRestClassifier wrapper into the same RF pipeline previously constructed, maintaining consistent hyperparameters and procedures. This OVR model achieved performance comparable to the original multiclass model, with a training accuracy of 1.0 and a testing accuracy of 0.9107 (Figure 5A). This similarity is expected, as both models share the same underlying architecture; the key distinction is that the OVR model trains a separate binary classifier for each class, whereas the original RF model predicts class probabilities using a single multiclass ensemble. In the OVR framework, each Wnt-Wls system is modeled independently against the remaining three. Permutation feature importance was then computed separately for each binary classifier, enabling evaluation of system-specific contributions from individual contact features. We highlight the results for Wnt3a and Wnt8a in Figure 5C-D, as both systems have been structurally characterized, providing experimentally validated contact information for comparison with our ML results. Feature importance results for Wnt1 and Wnt5a are provided in the Supplementary Material Figures S6-7. The selection criteria and structural sources for the four Wnt proteins are described in detail in the <u>Methods section</u>.

Overall, the highest feature importance scores observed in each binary classifier were substantially lower than those identified in the unified multiclass RF model. Notably, the contact pair I185-L297, which ranked as the most influential feature in the multiclass model, did not emerge as a top-ranking feature in any of the OVR classifiers. This suggests that certain evolutionary divergent positions, while informative for distinguishing among all four systems collectively, may contribute less in system-specific contexts where only a single class is being contrasted against the others.

In the case of Wnt8a (Figure 5C) the top-ranked contact pair is C319-I77. Among the other highly ranked pairs, most have not been directly investigated in existing structural studies of Wnt8a, with the exception of contacts involving residue C320, which ranked fourth, fifth, and sixth in importance. C320 lies immediately adjacent to C319 and together they form part of the conserved W-C-C motif. For Wnt3a (Figure 5C), the top-ranked contact pair is also C319-I77, reinforcing the importance of this interaction. These findings highlight the recurring role of the W-C-C motif in distinguishing system-specific binding signatures, even within independently trained binary classifiers.

As discussed in the previous section, C319 (corresponding to C334 in Wnt3a) lies at the center of the highly conserved W-C-C motif and plays a crucial role in interacting with the hydrophobic pocket of Wls. The second-ranked contact pair in Wnt3a is Q317-E86. Q317 aligns with W333 in Wnt3a, a residue shown in experimental studies to form extensive contacts with Wls^25^. Notably, substitution of W333 with alanine has been reported to abolish Wnt signaling activity^25^, highlighting its essential functional role. The consistent appearance of both C319 and C320 among the most predictive features for both Wnt3a and Wnt8a underscores the functional significance of the W-C-C motif. Given that this motif mediates critical contacts with the Wls hydrophobic pocket, our model’s identification of these residues as top-ranking features provides strong computational support for their biological relevance. Moreover, it suggests that the W-C-C motif may serve as a conserved structural element that enables functional divergence across Wnt isoforms, while maintaining a core binding mechanism essential for Wls recognition.

## 3. Discussion

In this study, we integrated atomistic MD simulations with ML to identify molecular determinants of the Wnt-Wls binding specificity across four members of the Wnt protein family and the Wls transmembrane protein. Understanding this many-to-one interaction is essential for explaining how Wnt signaling maintains fidelity across diverse biological contexts, and how mutations in either Wnt or Wls may drive signaling defects linked to diseases.

We focused on the following four biologically relevant Wnt proteins: Wnt3a and Wnt8a, which have available crystal structures, and Wnt1 and Wnt5a, based on scientific relevance (Figure 1A-C). MD simulations revealed a consistent dynamics pattern across systems: Wnt proteins exhibited high conformational flexibility, the Wls remained comparatively rigid in all complexes (Figure 1E). The RMSF analysis showed strong agreement between the two independent simulation replicas at the Wnt-Wls binding interface with differences limited to distal unstructured regions. Only two residues, R156 and T231 in Wnt1, showed replica-specific differences >1 Å near the interface, but neither impacted the downstream ML analysis: R156 was not among the 1,153 aligned contact pairs, and T231 had zero permutation feature importance. We therefore proceeded with Replica 1 for all ML analyses to ensure consistency while retaining relevant biophysical signals.

To enable ML-based analysis of Wnt-Wls interactions, we calculated all CL-CL distances between residues of Wnt and Wls proteins and defined a contact pair as any residue pair within 12 Å in at least one simulation frame. This threshold matches the long-range interaction cutoff used in our force field and aligns with prior work^29,30^. This approach yielded thousands of contact pairs per system: 2,208 for Wnt1-Wls, 2,019 for Wnt3a-Wls, 2,395 for Wnt5a-Wls, and 1,925 for Wnt8a-Wls (Figure 2C), providing a rich and consistent feature space across systems.

To enable cross-system comparison analysis despite differences in Wnt sequence length and composition, we developed a novel local structural alignment algorithm using RMSD as the primary metric. For each residue in Wnt1, Wnt3a, or Wnt5a, we extracted a 41-residue window (±20 residues) and aligned it to all 41-residue segments in Wnt8a, resulting in the lowest-RMSD match as the best structural counterpart. We set a 10 Å RMSD threshold determined through visual inspection of the simulations to retain structurally valid alignments, including those in flexible or disordered regions that would be excluded by stricter cutoffs (e.g., <2 Å). This method produced 1,153 structurally aligned Wnt-Wls contact pairs shared across all four systems. Unlike standard alignment approaches, such as that of Pavlova et al^30^, which prioritize amino type, our strategy emphasizes structural analogous residues. For example, T122 in Wnt8a was aligned to S145 in Wnt3a (RMSD = 3.75 Å), a mapping missed by multiple sequence alignment (MSA). In Pavlova et el.^30^ all peak residues differ in amino acid type between homologs, potentially reflecting either true mutational divergence or misalignment of structurally distinct residues. In contrast, several of our top-ranked contact residues are conserved in both structure and amino acid type, underscoring a key limitation of sequence-based methods and highlighting the generalizability and biological relevance of our RMSD-based alignment strategy.

To generate ML-ready input features, we extracted a time series of C_a_-C_a_ distances for the 1,153 shared contact pairs across all simulation frames. However, the fine-grained time resolution of MD simulations creates strong autocorrelation between consecutive frames, making naïve validation approaches such as random or K-fold splitting prone to severe data leakage. To address this, we developed an ACF-based data partitioning strategy. For each contact pair, we computed the ACF and defined the relaxation time (τ) the time at which the ACF decays to 0, as a temporal buffer to separate training and testing sets. This decorrelated split led to a modest but expected drop in test accuracy, reflecting improved robustness and a more realistic estimate of the model’s generalization performance.

To mitigate feature redundancy and multicollinearity, we applied a two-round hierarchical clustering strategy to the 1,153 aligned contact pairs. In the first round, contact features were grouped into 21 clusters based on structural proximity and functional annotation (Figure 4). A second round of subclustering yielded 200 clusters, from which we selected one representative feature per group. This step was critical to prevent overfitting, as many contact pairs in MD-derived contact maps often contain synchronized or spatially coupled features due to physical proximity. To assess the effectiveness of this feature reduction strategy, we calculated pairwise Pearson correlation coefficients and Kullback–Leibler (KL) divergence values across the selected 200 features. Only 3% of the feature pairs exceeded the 0.9 Pearson correlation threshold, a common cutoff for redundancy^30^, and just 8.8% showed KL divergence values below 0.4. These metrics confirm that our clustering pipeline produced a compact, decorrelated, and information-rich feature set, enhancing both model generalization and interpretability.

Using the curated set of 200 features, we trained an RF classifier that achieved 100% accuracy on the training set and 88.5% on the ACF-separated test set. This performance aligns with prior ML applications to MD trajectories^30^, where atom-level resolution and rigorous feature selection enable robust classification. Despite the high accuracy, permutation feature importance scores were modest, consistent with a distributed model in which no single feature dominates and classification depends on the collective contributions of many contact pairs across the Wnt-Wls interface. Among all features, the most predictive was the contact between I185 in Wnt8a and L297 in Wls. I185 is located adjacent to the conserved PAM site, a residue essential for binding, and aligns structurally (but not by amino acid identity) with M223 in Wnt1, L208 in Wnt3a, V243 in Wnt5a. This positional variability, despite a conserved binding core, may explain I185’s high predictive power. This finding mirrors principles observed in other systems such as the proline-rich motif in SH3 domain binding^27^, where conserved core residues mediate binding while flexible surrounding residues modulate specificity. We propose that I185 may function as a tunable determinant of Wnt-Wls binding specificity, modulating interactions or selectivity across isoforms.

Both the third- and fourth-ranked features involved C319, a conserved cysteine that forms the central residue of the W-C-C motif. Although this residue is conserved across all systems, appearing as C352 in Wnt1, C334 in Wnt3a, and C362 in Wnt5a, C319 consistently ranked among the top predictive features. This finding echoes prior work by Nygaard et al.^27^, which demonstrated that the W-C-C motif can undergo conformational switching and may act as a regulatory element in Wnt-Wls binding. Our results reinforce the view, showing that even fully conserved residues can function as critical determinants of binding specificity when they exhibit structural flexibility or context-specific behavior.

To gain further insight into the molecular mechanism underlying the protein binding interactions, we trained OVR binary classifiers for each Wnt protein. Unlikely the multiclass model, I185-related contacts were not top-ranked in any single OVR classifier, suggesting their value lies in differentiating between systems, rather than uniquely identifying any one. For Wnt3a, C319 again emerged as the top-ranked feature, followed by Q317-I77, which aligns with W333 in Wnt3a, a residue known to form critical contacts with Wls; mutation to alanine at this site abrogates signaling. For Wnt8a, C319 remains the strongest contributor, and C320, another member of the W-C-C motif, appeared in the next three highest-ranked contact pairs, further supporting the motif’s role in binding specificity. Altogether, our analysis highlights 16 contact pairs with high statistical significance and biological relevance. Of these, nine have been previously reported in Wnt-Wls structural studies, while the remaining seven represent potentially novel interactions not described in literature. Manual inspection confirmed that none of these novel contacts have been annotated, making them compelling targets for future experimental validation. These findings highlight the power of simulation-guided ML to uncover distributed specificity mechanisms in protein-protein interfaces and point toward several promising targets for future validation.

## 4. Conclusions

We uncovered key structural features that govern Wnt-Wls binding specificity using a novel integration of long time-scale MD simulations and interpretable ML. Our RMSD-based alignment strategy enabled residue-level mapping across divergent Wnt proteins, revealing contact pairs conserved in position but variable in identity. By applying ACF-informed data split and hierarchical clustering, we built compact, generalizable models that achieved high predictive accuracy without overfitting. Importantly, our analysis identified 16 contact pairs with high statistical and biological relevance, seven of which have not been reported previously. These include I185 (adjacent to the PAM site) and multiple residues in the W-C-C motif which collectively appear to modulate binding specificity. These findings demonstrate that specificity is encoded not by a single residue, but by a distributed set of context-sensitive contacts. Ur approach offers a generalizable framework for dissecting protein-protein interactions and can inform mutagenesis experiments or therapeutic targeting of structurally conserved yet functionally tunable interfaces.

## 5. Methods

### 5.1 Molecular Dynamics Simulations

#### System Selection

For this study, we identified Wnt1-, Wnt3a-, Wnt5a- and Wnt8a-Wls systems based primarily on the availability of their experimental structures and their scientific relevance, confirmed by a bibliometric analysis. For the bibliometric analysis, we used publication metrics extracted from the Web of Science Core Collection database and considered the total number of publications, citations, and H-index. Our protein selections combined experimental structural data availability with current bibliometric indicators of research significance and relevance.

#### System Preparation

We used the same protocol described herein to set up and run all the simulations performed in this work. All systems were prepared with CHARMM-GUI^45–47^ using the membrane builder module containing human Wnt-Wls complex, a membrane bilayer composed of 100% palmitoyloleoylphosphatidylcholine (POPC), and TIP3 water^48^. Each system was simulated in a tetragonal cell with approximate dimensions of 110×110×160 Å^3^. The corresponding lipid counts for the upper and lower leaflets, along with the total number of atoms are summarized in Table 1. We built four different Wnt-Wls systems using Wnt1, Wnt3a, Wnt5a, and Wnt8a. To generate the initial coordinates of Wnt3a-Wls and Wnt8a-Wls complexes we used the experimental structures PBD ID 7DRT^25^ and 7KC4^27^, respectively. For the Wnt1- and Wnt5a-Wls models, we used Phyre2.2^42^ (see below for more modeling details). Ions (Na^+^ and Cl^-^) were added to neutralize each system at a concentration of 0.15 M. The CHARMM ACE and CNEU terminal groups were applied for both Wnt and Wls chains to neutralize both ends of the proteins.

**Table 1.** Simulation details for each Wnt-Wls system.

| Wnt-Wls System | Atom number | Lipid number<br>(top/bottom leaflet) | Replica | Simulation length (μs) |
| --- | --- | --- | --- | --- |
| Wnt1-Wls | 182,554 | 155/158 | 2 | 1.5 |
| Wnt3a-Wls | 180,086 | 155/158 | 2 | 1.5 |
| Wnt5a-Wls | 179,768 | 155/158 | 2 | 1.5 |
| Wnt8a-Wls | 175,583 | 155/158 | 2 | 1.5 |

#### Simulation Protocol

Using NAMD 2.14^49^, all systems followed the minimization and equilibration protocol provided by CHARMM-GUI^46^. The protocol included 6 steps of equilibration, where components initially restrained were progressively relaxed at each stage of the equilibration process. NAMD 2.14 was used in this phase of the simulation set up, since this version implemented the collective variable module required for the restrained equilibration. The production run simulations were performed using NAMD 3.0 on the AiMOS supercomputer, which provided a calculation speed-up compared to NAMD 2. The production runs were performed in the *NPT* (constant number of particle *N*, pressure *P*, and temperature *T*) ensemble, at the constant temperature of 303.15 K and pressure of 1 atm. The temperature was maintained using Langevin dynamics, with a damping coefficient of 0.5 ps^-1^. Constant pressure was maintained by the Langevin piston Nose-Hoover method^50^, with a decay period of 50 fs and piston period of 100 fs. The non-bonded cutoff distance for short-range interactions was 12 Å, with switching at 10 Å. A 1-4 scaling factor of 1 was used with a switch distance of 10 Å. The particle mesh Ewald (PME) method^51^, with a 1 Å grid spacing, was used for long-range electrostatics. Hydrogen atom bond lengths were restrained with SETTLE^52^ to allow an integration step of 2 fs. Frames were saved every 20 ps to be used for analysis. All systems were subjected to periodic boundary conditions using a tetragonal box. Table 1 provides a summary of the simulations performed in this work. We constructed and ran two independent replicas per system.

#### Serine-Palmitoleate Modeling Approach

Wnt biology involves a mono-unsaturated palmitoleate (PAM) covalently attached to a conserved serine residue in all human Wnts^25,27^, however the CHARMM36 force field parameters necessary to simulate this lipidated serine were unavailable at the moment of starting this work. Generating new forcefield parameters would have required quantum mechanical calculations that were outside the scope of this study. We then adopted an alternative approach and generated the parameter files leveraging the existing ones for molecules with similar atom types. We determined this parameterization strategy sufficient for the goal of our work which was focused on the Wnt-Wls interactions rather than protein-membrane interactions. Nonetheless, our strategy captures PAM’s essential function, such as its anchoring to the lipid bilayer^27^.

Because of the chemical features resembling Serine-PAM (SERP), we first examined the parametrization of the cysteine-palmitate residue (CYSP), i.e. a lipidated cysteine. We then analyzed the correspondence between CYSP forcefield parameters to existing small molecules and applied the same chemical mapping to derive SERP parameters. For instance, if we consider the CYSP sidechain and its angle parameter composed of cysteine’s sulfide functional group with a lipid attached, we found that one angle at this interface is formed by cysteine’s Cβ, Sγ, and palmitate’s C1. The parameters describing the angle of these 3 atom types (CT2-S-CL) are equivalent to those of ethyl methyl sulfide (CG321-SG311-CG331). Using this approach of finding equivalent atom types in corresponding small molecules, we derived the bond, angle, and dihedral parameters for the interface between the gamma oxygen of serine and its attached palmitoleate lipid (Supplementary Material, Table S2). PAM aliphatic carbon chain parameters were directly adopted from di-palmitoleic-phosphatidylcholine in the CHARMM36 force field, as its lipid tails contained the same chemical features as SERP, i.e. 16 carbon atoms and a C9-C10 cis double bond.

#### Phyre2 Homology Modeling of WNT1 and WNT5a

Since the experimental structures of Wnt1 and Wnt5a have not yet been resolved, we modeled them using the Phyre2.2^42^. Although AlphaFold^53^ also generates highly accurate protein structures, we specifically selected Phyre2.2 for its one-to-one threading capability, allowing explicit use of experimentally determined structural templates. This approach enabled precise modeling of biologically critical features, such as the accurate positioning and orientation of the conserved PAM residues interacting with the Wls hydrophobic tunnel between TM4 and TM5.

Our rationale for using Wnt3a as a template was due to the high-resolution structure of 2.2 Å and the high sequence identity (46%) between Wnt1, Wnt5a and Wnt3a. The input sequences of Wnt1 and Wnt5a were derived from UniProt with UniProt IDs P04628 and P41221 respectively. The Phyre2 output model of Wnt1 contains residues between 32 and 369 and that of Wnt5a residues between 32 and 370. Because for Wnt1 the final C-term leucine (L370) was not modeled from Phyre2, we decided to exclude it from our simulations. This residue is situated away from the Wnt-Wls interface and therefore not involved in protein-protein interactions.

#### Disulfide Bond Modeling of Phyre2.2 output

Human Wnts consist of 23 or 24 conserved cysteine residues arranged with similar spacing, indicating that protein fold relies on several disulfide bonds^54^. A prior biochemical study supported this hypothesis by showing how the mutation of Wnt3a disulfide bonding cysteines diminished the signaling capabilities^55^. We incorporated these essential disulfide bonds into the models of Wnt3a and Wnt5a using the Wnt3a experimental structure (PDB ID: 7DRT) as a reference. We rationalized that Wnt3a would be a reasonable template due to its high sequence identity and sufficient structural resolution to resolve disulfide bonds. Thus, any disulfide bonds dissimilar to 7DRT were excluded from the final model. The summary of these disulfide bonds is shown in Supplementary Material, Table S3.

#### Glycan Modeling

For Wnt8a, two glycans consistent with the cryo-EM structure (PDB ID: 7KC4) were incorporated into the simulation. One is a 6-mer, N-linked oligosaccharide at N103, which was shown with mutagenesis experiments to have an important functional role^27^ as it also interacted at the Wnt8a-Wls interface. The second glycan is a 2-mer, situated away from the Wnt8a-Wls interface. We did not model the glycans for Wnt1, Wnt3a, and Wnt5a. The Wnt3a crystal structure (PDB ID: 7DRT) contains 1 glycan at N87 and was excluded because its short chain (2-mer) appears to not interact with Wls. To the best of our knowledge, no reports are published for Wnt1 and Wnt5a that determine the exact glycan chain lengths. Moreover, based on a visual inspection of the proposed N-linked glycosylation sites in the UniProt database, we determined that these glycans would not contribute to stabilizing the Wnt-Wls interface. In particular, the Asn residues were situated too far, with their side-chain orientation directed away from Wls.

#### Analysis of the Simulation Data

The RMSD and RMSF of the Wnt and Wls proteins reported in this study were calculated using ptraj^56^. We calculated the RMSDs for CL atoms after performing a protein backbone alignment of structures to the first frame of the production run. We performed a rolling window average of 10 frames (200 ps) to identify better for convergence and structural stability of the systems. We calculated the RMSF for each residue using the block averaging method. The last 500 ns of 1.5 µs simulations were extracted for each replicate and split into 5 blocks (100 ns per block). The averages and standard deviation were then calculated from these trajectory subsets. All RMSF calculations were performed after aligning the protein backbone atoms to the first frame of the production run.

#### Contact Pairs estimation and Local Structural Alignment algorithm

We estimated the contact pairs for the four systems under investigation using a method based on cutoff distance between the CL atoms of the residues belonging to the different proteins. For each system we calculated the distance between the CL atoms of one protein and those of the other protein at each simulation time step. If the CL-CL distance fell below 12 Å at least in one frame, then the residues to which the CL atoms belong to were considered as contact pair.

Following this approach, the numbers of contact pairs result to be 2,208 for Wnt1-Wls, 2,019 for Wnt3a-Wls, 2,395 for Wnt5a-Wls, and 1,924 for Wnt8a-Wls, as illustrated in Figure 2C. This cutoff-distance based method does not allow to extract the common contact pairs shared by the four Wnt-Wls systems because the Wnt proteins are characterized by different sequence length and amino acid composition.

For comparing the different Wnt sequences, we implemented a RMSD-based structural alignment which offers a more robust method for identifying conserved Wnt-Wls contacts across all four Wnt proteins (Figure 2A). We chose Wnt8a as reference system because it has the longer sequence of the two available crystallographic resolved structure (at the time of analysis) and aligned Wnt1/3a/5a to Wnt8a. To identify local structural correspondences between the different sequences, we then conducted a search for each amino acid *i* in Wnt1, Wnt3a, and Wnt5a. The local structure around each *i* was defined as a 41-amino-acid segment, spanning residues [*i-20, i+20*]. This range was selected to ensure coverage of the longest secondary structural elements found in Wnt proteins, such as alpha-helices (∼25 amino acids) and beta-sheets (∼34 amino acids). The chosen range ensures that our procedure captures the entire secondary structures each time, minimizing the risk of identifying multiple overlapping matching regions within the same structure, or structures with similar features, such as alpha-helices. For each *i*, a corresponding match was sought in Wnt8a by iterating through all possible 41-amino-acid segments centered at residue *j* in Wnt8a. The residue *j* that provided the best fit for *i* was identified based on the minimum value of the RMSD between the two segments, [*i*-20, *i*+20] in Wnt1/3a/5a and [*j*-20, *j*+20] in Wnt8a.

The RMSD was calculated in two steps. First, an initial RMSD alignment was performed on the two 41-amino-acid segments, with the conserved Wls protein included in the alignment. Including the Wls in this step ensured optimal global alignment, as all systems share the same Wls structure. Subsequently, the RMSD was recalculated, focusing exclusively on the two 41-amino-acid segments. The best structural match for each residue *i* in Wnt1/3a/5a corresponds to the residue *j\** in Wnt8a that minimizes the RMSD between segments [*i*-20, *i*+20] and [*j*\*-20, *j*\*+20]. This two-step approach ensured both the alignment quality and the reliable identification of local structural correspondences between the Wnt proteins. For residues located at the terminal ends, the segment range was adjusted to [*i*, *i*+40] for the N-terminal and [*i*−40, *i*] for the C-terminal. The range for *j* is also adjusted in the same manner.

Mutations such as insertions and deletions alter protein length, preventing every residue *i* in Wnt1/3a/5a from having a single, exact match in Wnt8a. Consequently, some residues may lack corresponding matches altogether. However, our RMSD method consistently identifies the best match for each residue, defined as the pair with the smallest RMSD value, even in cases where residues lack a suitable match in Wnt8a. We implemented a cutoff value following the two-step RMSD procedure to filter out poorly fitting residue pairs while conserving high-quality matches. The cutoff value was determined empirically, as no prior knowledge exists to define a universal threshold. To establish an appropriate value, we analyzed the distribution of all minimum RMSD values and visually inspected the quality of local structural alignment of residue pairs corresponding to these values (Figure 2B). This analysis revealed that pairs with RMSD values exceeding 12 Å consistently represented poor fits and were excluded. Residue pairs with RMSD values in the range of 10-12 Å constituted a “gray area,” often corresponding to amino acids located at the termini of well-folded secondary structures or at boundaries of disordered regions. To optimize filtering, a cutoff value closer to 10 Å was recommended for strict selection, while a more inclusive approach allowed for values up to 12 Å. This range provides flexibility, enabling the adjustment of the threshold based on the desired balance between specificity and inclusivity in the identification of high-quality residue matches. We used 10 Å as a cutoff for our system. We repeated this process for all possible amino acid ranges on Wnt1/3a/5a to determine a mapping of residue indices to Wnt8s, ensuring consistent amino acid numbering between each of the 4 systems.

### 5.2 Machine Learning Protocol

Figure 3 provides a high-level overview of the data preparation and machine learning pipeline, each of which will be described in subsequent sections.

#### Data Preparation

The result of the MD simulations is a time series where each frame contains the Cartesian coordinates of all the atoms of the system. We used the following procedure to convert the data into a training set for our machine learning workflow. To identify the amino acids involved in the protein-protein interactions and build an interaction matrix, we define a contact pair between Wnt and Wls when the distance between the Wnt and Wls C_a_ atoms happens to be within 12 Å at least for a single frame during the whole simulation length. Compared to prior literature^29,30^, we selected the value of 12 Å because it corresponds to the non-bonded cutoff distance for short-range interactions used for the simulations. We recorded the distances every 20 ps to up to 1500 ns, i.e. 75000 frames were analyzed and preprocessed.

#### Training and Testing Splits

The ACF of contact pair distances was computed using in-house code reported in our GitHub repository. To ensure a consistent estimation of error bars across the time interval t, the same sliding window approach previously employed for calculating the orientational autocorrelation function was applied during the ACF computation^57^. The resulting ACF profiles were then used to identify optimal frames within each Wnt trajectory for generating the training and testing splits. The ACF for each of the four systems averaged over 75000 frames, all 1153 contact pair distances. ^58^. For training and testing splitting, we defined the splitting time point τ as the value at which the ACF decays to zero, in this way we determined the time gap for which two time points are not correlated anymore. We identify τ as the split point between the training and testing data set and consider the testing data set the data between 0 and τ. To ensure that the two sets are completely uncorrelated, we excluded the data from τ and 2xτ, and used the data between 2xτ and the end of the simulations as the training set. This τ value was also used as the gap parameter in the splitting strategy for both the learning curve analysis and cross-validation, as discussed in subsequent sections.

#### Feature Selection

Given the high dimensionality of the contact pair features and the presence of strong correlations among them, we applied a two-round hierarchical clustering strategy to group similar or correlated contact pairs and select a representative from each cluster. This approach was employed to reduce redundancy, enhance model generalization, and reduce the risk of overfitting. Both rounds of clustering were performed using scikit-learn AgglomerativeClustering module^59^. Silhouette Scores (SS) were used to evaluate the clustering fit for both rounds. In the first round, clustering was performed on the contact index pairs using Euclidean distance as the metric and Ward’s linkage method. In the second round, the inverse of the Spearman correlation coefficient was used to transform the contact pair distances into a precomputed distance matrix suitable for clustering.

After clusters were identified from the first round, representative contact pairs were selected based on the following procedure: (1) If the cluster contained more than four contacts pairs, we applied a second round clustering to the cluster and one contact pair was randomly sampled (random seed = 0) from each of the subsequent subclusters; (2) if the cluster contained two to three contacts, one contact pair was originally sampled; and (3) if the cluster contained a single contact pair it was used. This selection procedure resulted in a final set of 200 contact pairs, which were used as input features for downstream analysis.

#### Building a Random Forest Machine Learning Classifier to Identify Important Features

We performed a Random Forest multiclass classification using Scikit-Learn, with accuracy as the scoring metric and Gini impurity as the splitting criterion. Hyperparameter optimization was conducted using grid search with three iterations of 10-fold cross-validation (random seeds = 9, 47, 512). Since the input features exhibit temporal dependencies, we employed the TimeSeriesSplit cross-validator from scikit-learn to ensure proper data partitioning in the context of time series samples. The largest τ value obtained from the ACF analysis across the four systems was used as the gap parameter in the TimeSeriesSplit. The optimized parameters included the number of trees (n_estimators = 5, 10, 25, 50, 200, 1000), minimum samples per leaf (min_samples_leaf = 1, 1641, 4104), and tree depth (max_depth = None, 1, 3, 5, 10). The values for min_samples_leaf were calculated by scaling the training set by 0.1 and 0.25, then dividing by the number of folds (10). Specifically, train_size*(0.1/10)=1,641 and train_size*(0.25/10)=4,104. All other parameters were kept at default settings specified by Scikit-Learn.

The best model was selected based on average accuracy across iterations and folds. Several parameter combinations yielded perfect accuracy (1.0) on both training and testing datasets. Among these, we selected the configuration with the highest min_samples_leaf, the lowest max_depth, and the lowest n_estimators to favor model simplicity and reduce the risk of overfitting. Final model hyperparameters (n_estimators = 200; max_depth = 3; min_samples_leaf = 1641) were validated using a learning curve generated with a random seed of 9 and 20-fold TimeSeriesSplit, evaluated as a function of the percentage of training data. To identify the features that differentiated Wnt systems, we computed permutation feature importance on the test set, running 100 iterations with the final model. Using the test set ensures a robust assessment of feature importance, reflecting the trained model’s performance on unseen data. The features with the highest permutation importance were examined and compared against findings reported in the experimental literature. We also applied a one-vs-the-rest classification strategy using the OneVsRestClassifier from scikit-learn. The same permutation feature importance analysis was then performed to identify features that are critical for distinguishing each individual Wnt system.

## Declaration of competing interests

The authors declare no competing interests.

## Code Availability

All scripts and codes used for local structure alignment, data analysis, and machine learning model development are available at https://github.com/CCCofficial/MDML-WntWls-Protein_Interactions.git.

## Author contributions

KJC, TJC, TVP, and SC conceived the study. KJC selected the systems, set up and carried out the atomistic simulations. TJC designed the machine learning framework. JS analyzed the simulations and carried out the machine learning analysis. MS developed the training/testing splitting dataset method based on autocorrelation functions. TJC and JS supervised MS work. SC and TJC provided guidance and supervision. SC, KJC, TJC, JS wrote the paper. All authors edited the paper.

## Acknowledgments

This material is based upon work supported by the National Science Foundation under Grant No. DBI-1548297. We acknowledge computational time in the School of Chemical Sciences Computer Center at University of Illinois Urbana-Champaign. We thank the computational support from the Artificial Intelligence Multiprocessing Optimized Systems (AiMOS) made available by the IBM Research AI Hardware Center and Rensselaer Polytechnic Institute’s Center for Computational Innovation. We thank Prof. Laura Burrus, Prof. Marc O Anderson, and Lisa Galli from San Francisco State University for the fruitful scientific discussions.

## Notes

### Competing Interest Statement

The authors have declared no competing interest.

